# Synthetic amyloid beta does not induce a robust transcriptional response in innate immune cell culture systems

**DOI:** 10.1101/2021.09.14.460110

**Authors:** I.Y. Quiroga, A.E. Cruikshank, K. S. M. Reed, M.L. Bond, B.A. Evangelista, J.H. Tseng, J. V. Ragusa, R. B. Meeker, H. Won, S. Cohen, T.J. Cohen, D.H. Phanstiel

**Author notes:** Equal contribution (I.Y. Quiroga, A.E. Cruikshank).

## Abstract

Alzheimer’s disease (AD) is a progressive neurodegenerative disease that impacts nearly 400 million people worldwide. The accumulation of amyloid beta (Aβ) in the brain has historically been associated with AD, and recent evidence suggests that neuroinflammation plays a central role in its origin and progression. These observations have given rise to the theory that Aβ is the primary trigger of AD, and induces proinflammatory activation of immune brain cells (i.e. microglia), which culminates in neuronal damage and cognitive decline. In order to test this hypothesis, many in vitro systems have been established to study Aβ-mediated activation of innate immune cells. Nevertheless, the transcriptional resemblance of these models to the microglia in the AD brain has never been comprehensively studied on a genome-wide scale. To address this, we used bulk RNA-seq to assess the transcriptional differences between in vitro cell types used to model neuroinflammation in AD, including several established, primary and iPSC-derived immune cell lines (macrophages, microglia and astrocytes) and their similarities to primary cells in the AD brain. We then analyzed the transcriptional response of these innate immune cells to synthetic Aβ. We found that human induced pluripotent stem cell (hIPSC)-derived microglia (IMGL) are the in vitro cell model that best resembles primary microglia. Surprisingly, synthetic Aβ does not trigger a robust transcriptional response in any of the cellular models analyzed, despite testing a wide variety of Aβ formulations, concentrations, and treatment conditions. Finally, we found that bacterial LPS and INFγ activate microglia and induce transcriptional changes similar to those observed in disease associated microglia present in the AD brain, suggesting the potential suitability of this model to study AD-related neuroinflammation.

---

**A**lzheimer’s disease (AD) affects about 5.8 million people in the USA and nearly 400 million people worldwide. The estimated cost for health care in the United States is 301 billion dollars and this number is expected to increase to 1.1 trillion dollars by 2050 when AD is projected to affect 13.8 million people^1^. In order to improve diagnosis and treatment options for AD it is imperative to have a better understanding of the molecular mechanisms underlying AD.

The accumulation of Amyloid beta (Aβ) in the brain has been associated with AD since the discovery of this pathology in 1906. This correlation has been supported by a number of findings, including the identification of genetic mutations causing misregulation of Aβ production and processing as the cause for familiar AD^2^ and the evidence that identifies Aβ deposition upstream of Tau tangle formation and neuronal death^3–8^. As a result, Aβ deposition has been proposed as the main cause for the development of AD, a theory known as the amyloid cascade hypothesis. Nevertheless, recent findings have raised questions about this hypothesis, including the failure of drug trials targeting Aβ accumulation and the observation of similar patterns of Aβ deposition in AD and healthy brains^9^. This evidence suggests that although closely associated, Aβ accumulation might not be the only causal factor for the development of AD and other elements might contribute to a more complex etiology.

Evidence pointing to chronic neuroinflammation as a crucial contributor to AD progression has accumulated over the last decade. Patients with AD have higher levels of neuroinflammation when analysing inflammatory markers in postmortem AD brains or live neuroimaging^10–15^. Resident brain immune cells from patients with AD express more genes associated with immune pathways and phagocytosis than controls indicating a state of chronic activation^16^. Finally, recent GWAS studies revealed enrichment of disease-associated variants at monocyte, macrophage, and microglia regulatory loci implicating innate immune cells as key mediators of AD risk^17–19^. This evidence has led to the hypothesis that in the Alzheimer’s brain, Aβ triggers an exacerbated proinflammatory activation of brain innate immune cells, particularly microglia and to a lesser extent astrocytes, resulting in a chronic neuroinflammatory state that causes neuronal damage and cognitive decline.

Several in vitro models have been established to further investigate the role of immune cells in AD and the functional consequences of AD risk genes in these cells. Many of these models involve the exposure of mice or human innate immune cell types—either primary cells, established cell lines or human induced pluripotent stem cell (hIPSC)-derived microglia (IMGLs) to synthetic Aβ. It has been assumed that these in vitro scenarios would trigger an inflammatory response similar to the one observed in the human AD brain, but to the best of our knowledge comprehensive profiles of the transcriptional response to this stimulus in vitro have not been reported. Here we assess the transcriptional differences between in vitro cell types broadly used to model neuroinflammation— including THP-1 macrophages, human peripheral blood mononuclear cell derived macrophages (PBMC Mϕ), U-87 Astrocytes, HMC3 microglia-like cells, and IMGLs—and study their transcriptional response to synthetic Aβ. We found that IMGLs are the in vitro cell model that best resembles primary microglia, but that synthetic Aβ does not trigger a robust transcriptional response in any of the cells analyzed. Finally we found that treatment of IMGLs with bacterial LPS/INFγ induces transcriptional changes similar to those observed in disease associated microglia present in the AD brain, suggesting the potential suitability of this model to study AD-related neuroinflammation.

## Results

### Transcriptional comparison of cultured cell types to study AD

In order to determine which of the currently available and most widely used in vitro immune models best resembles the transcriptional signature of human microglia, we performed RNA-seq in PBMC Mϕ, THP-1 macrophages, HMC3 microglia, U-87 astrocytoma cells, and IMGLs. We used principal component analysis (PCA) to compare the transcriptomic profiles of these cells with already published data from primary human adult and fetal microglia as well as with other related immune cell types^20^. Our results showed that IMGLs cluster closely to both human adult and fetal microglia and further away from other primary immune cells such as monocytes and dendritic cells (**Figure 1A**). PBMC Mϕ and THP-1 macrophages are the next closest in proximity while the HMC3 microglia established cell line has the least similarities to primary microglia and, surprisingly, clusters almost perfectly with the U-87 astrocytoma cell line. We used variance partitioning to determine how much of this clustering could be driven by the fact that the data from these cell types was generated by two separate labs (variancePartition^21^). We calculated the percentage of variance explained by cell type, lab of origin, biological replicate, and technical replicate and found that on average 71% of variance was due to differences in cell type, while only 12% was attributable to lab of origin. Investigation of microglia marker genes in each of these cell types further confirmed the similarity between IMGLs and primary microglia **(Figure 1B)**. IMGLs showed high expression of TREM2, P2RY12, AIF1 (IBA1) and PLCG1, similar to primary adult and fetal microglia. THP-1 macrophages exhibited the next most similar pattern of expression at most but not all genes, while the HMC3 microglia established cell line exhibited low expression for the majority of these microglia marker genes. These results suggest that, among the cells investigated, IMGLs best resemble primary human microglia and are a suitable in vitro model to study neuroinflammation.

**Fig. 1.**
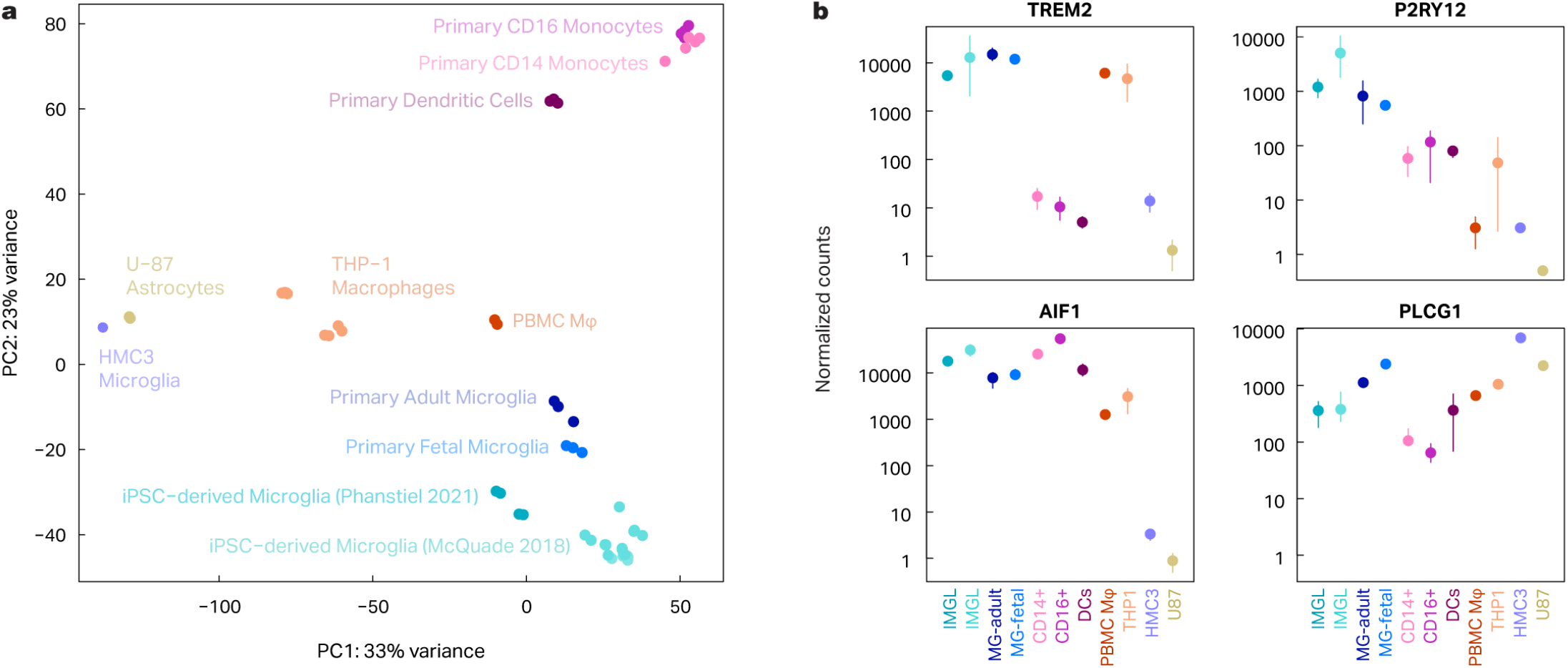
IMGL are the in vitro cell cultures that best resemble human microglia. Bulk RNA-seq was performed in different innate immune in vitro cell cultures and compared to already published data from related cell types. **(a)** Principal component analysis demonstrates that IMGL (light and dark turquoise) is the cell type that clusters most closely to cultured human fetal and adult microglia (dark and light blue). Additionally, these cells are distinct from human CD14+ monocytes (light pink) and CD16+ inflammatory monocytes (medium purple), and dendritic cells (dark purple). PBMC Mφ (red) and THP-1 cells (orange) are next in similarity and surprisingly the microglia HMC3 cell line (light purple) clusters more closely to the astrocytoma U-87 cell line (mustard) than to any microglia cell type analyzed. **(b)** The analysis of normalized counts of a subset of microglia markers show similar trends.

### Synthetic Aβ does not induce a robust transcriptional response

We next sought to characterize the global transcriptional response of IMGLs to synthetic Aβ to model the changes that occur in microglia in the AD brain. We treated IMGLs with 1 μM oligomeric Aβ for 24 hours and performed stranded, paired-end, total RNA-seq. Surprisingly, we observed no significant changes in gene expression in response to Aβ treatment (DESeq2, p < 0.01, absolute fold change > 2) (**Figure 2**). To explore the degree to which treatment conditions affect these results, we turned to THP-1 cells which are a more tractable system and express many microglia marker genes such as TREM2, a cell surface receptor known to bind Aβ^22^.

**Fig. 2.**
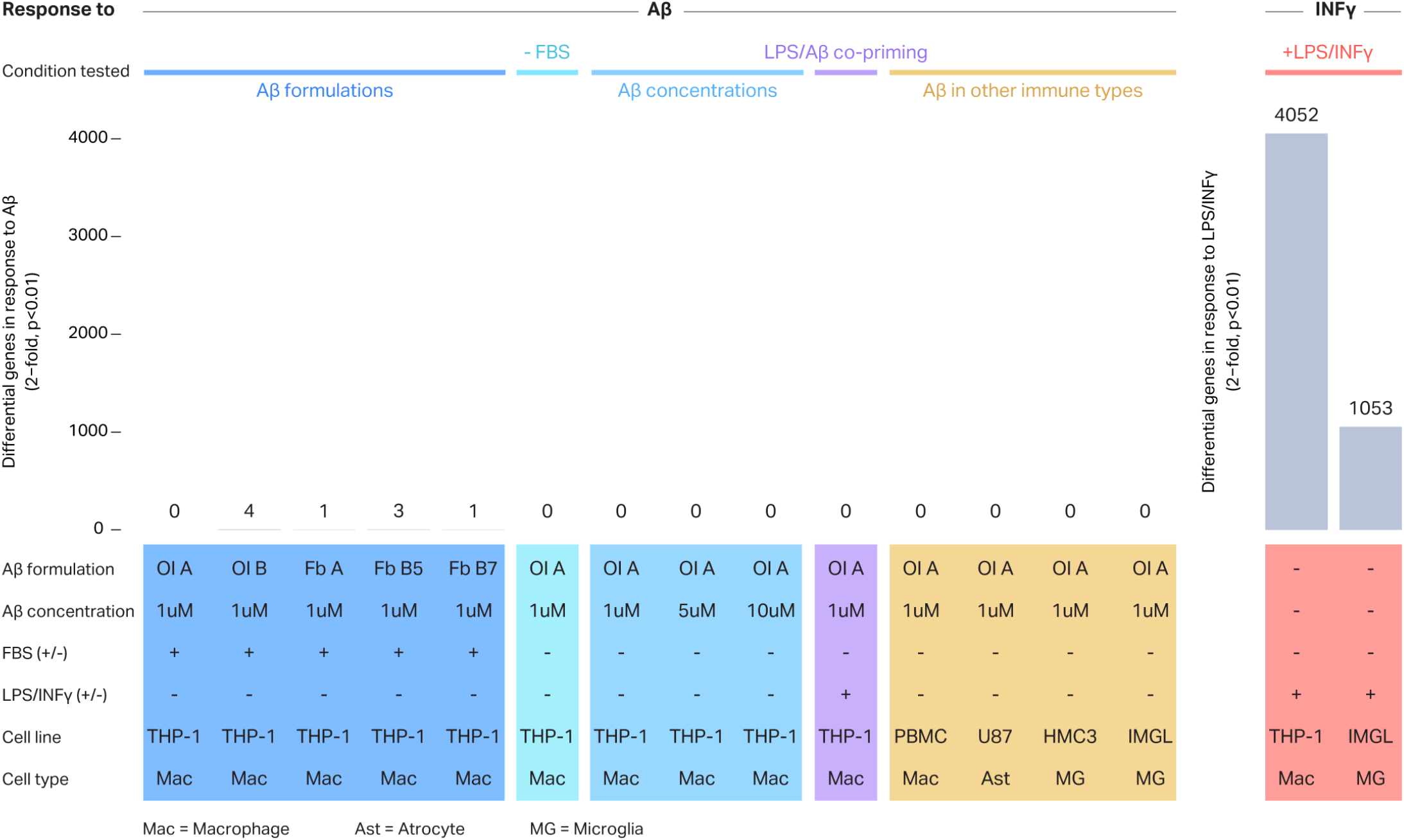
Transcriptional response of immune in vitro cell cultures exposed to different proinflammatory stimuli. Bulk RNA-seq was performed in different innate immune in vitro cell cultures exposed to proinflammatory stimuli during 24h. THP-1 cells were treated with different Aβ formulations including Aβ oligomers (Ol) and Aβ fibrils (Fb) obtained using different protocols (dark blue); Aβ in the absence of FBS (turquoise), with different Aβ concentrations (light blue), or co-priming using LPS/ INFγ and Aβ together (purple). Additionally, other immune cell types were treated with Aβ (orange). Lastly THP-1 cells and IMGLs were treated with 10nM LPS+20nM INFγ (red). Differential genes in response to Aβ or LPS/ INFγ were determined by comparing to condition-matched controls using DESeq2, p< 0.01, absolute fold change > 2.

We prepared synthetic Aβ in the form of oligomers or fibrils following two different previously published protocols: protocol A described by Tseng et al^23^ or protocol B described by Abud et al^24^. Successful generation of oligomers and fibrils was confirmed by western-blot (**Figure 3A**). We also confirmed that both THP-1 cells and IMGLs are capable of phagocytosing synthetic Aβ after 1h of incubation (**Figure 3B, Supplementary Figure 1**).

**Fig. 3.**
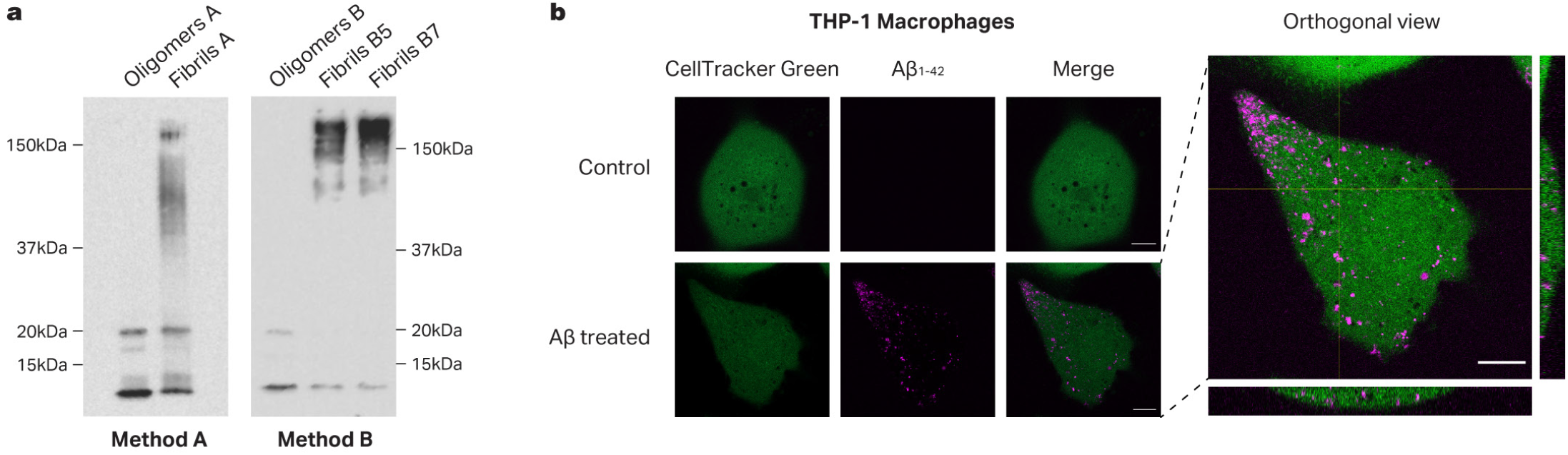
Aβ is incorporated into THP-1 cells. **(a)** Aβ Oligomers and fibrils were generated using either Tseng et al, 2017 (Method a) or Abud et al. 2017 (Method b) protocol. In the last case, fibrils were generated during either 5 or 7 days of incubation (B5 and B7, respectively). 120 ng of Aβ were run in an electrophoresis on a 12% SDS-PAGE followed by western blot using anti-Aβ primary antibody (6E10 Biolegend). **(b)** THP-1 cells were labeled with CellTracker Green CMFDA dye and incubated for 1h with 2μg/mL of oligomeric β-Amyloid_1-42_-HiLyte Fluor 555. Confocal images were acquired on an inverted Zeiss 800 laser scanning confocal microscope. Scale bar, 10 μm.

Next, we proceeded to treat THP-1 cells with Aβ under various conditions and performed RNA-seq to analyze the global transcriptional response. As observed previously in IMGLs, when THP-1 macrophages were treated with the different Aβ formulations (**Figure 2**, dark blue), no robust transcriptional response was observed in any condition. From this point forward, we utilized the oligomeric form of Aβ since it has been previously described as the more “active” form of Aβ^25–27^. We next eliminated the presence of fetal bovine serum (FBS) during the treatment to determine whether it was interfering with the Aβ incorporation, since it has been suggested that the albumin present in the FBS could act by “sequestering” the soluble Aβ in the media, reducing the availability of Aβ capable of binding the cellular receptors^28^. No significant transcriptional response was seen in the absence of FBS during the Aβ treatment. We then decided to test different concentrations of Aβ. Again, no response was seen with any of the Aβ concentrations tested ranging from 1 to 10μM. Following the relatively new hypothesis that microbial presence in the brain could be a contributing factor to the development of AD in addition to Aβ accumulation^29–32^, we decided to test whether Aβ has any synergistic effect when the cells were co-primed with LPS and INFγ. We found no significant differences when treating THP-1 cells with this combination compared to LPS/INFγ alone. Additionally, we tested whether Aβ had a significant effect in any of the other innate immune cell cultures used previously. No significant transcriptional response to Aβ was seen in either PBMC Mϕ, U-87, or HMC3 cells. As an alternative method to activate the inflammasome and also as a positive control for proinflammatory activation, we treated THP-1 cells with LPS and INFγ. As expected, THP-1 cells treated with this stimulus displayed a robust transcriptional response, (DESeq2, 4053 differential genes, p < 0.01, absolute fold change > 2) (**Figure 3**) Finally, we treated IMGL with LPS and INFγ and observed a robust transcriptional response (DESeq2, 1053 differential genes, p < 0.01, absolute fold change > 2) although smaller to the one observed in THP-1 macrophages.

### LPS/INFγ activation of IMGLs mirrors transcriptional changes seen in DAM

In contrast to amyloid beta, and in agreement with a recent study^33^, LPS/INFγ treatment induced a robust transcriptional response in both THP-1 macrophages and IMGLs (**Figure 4A-B**). Upregulated genes included key proinflammatory genes IL1B, CD38, and CCL2 and were enriched for classical inflammation pathways including NF-κB and Toll-Like Receptor Signaling (Homer, p < 0.05). Because many of these genes and pathways are upregulated in AD microglia and because AD pathogenesis has been associated with viral and bacterial infection^32,34,35^, some groups have used LPS or LPS/INFγ to stimulate innate immune cells either in vitro or by direct injection in animal’s brains as a model of AD-associated neuroinflammation^36,37^.

**Fig. 4.**
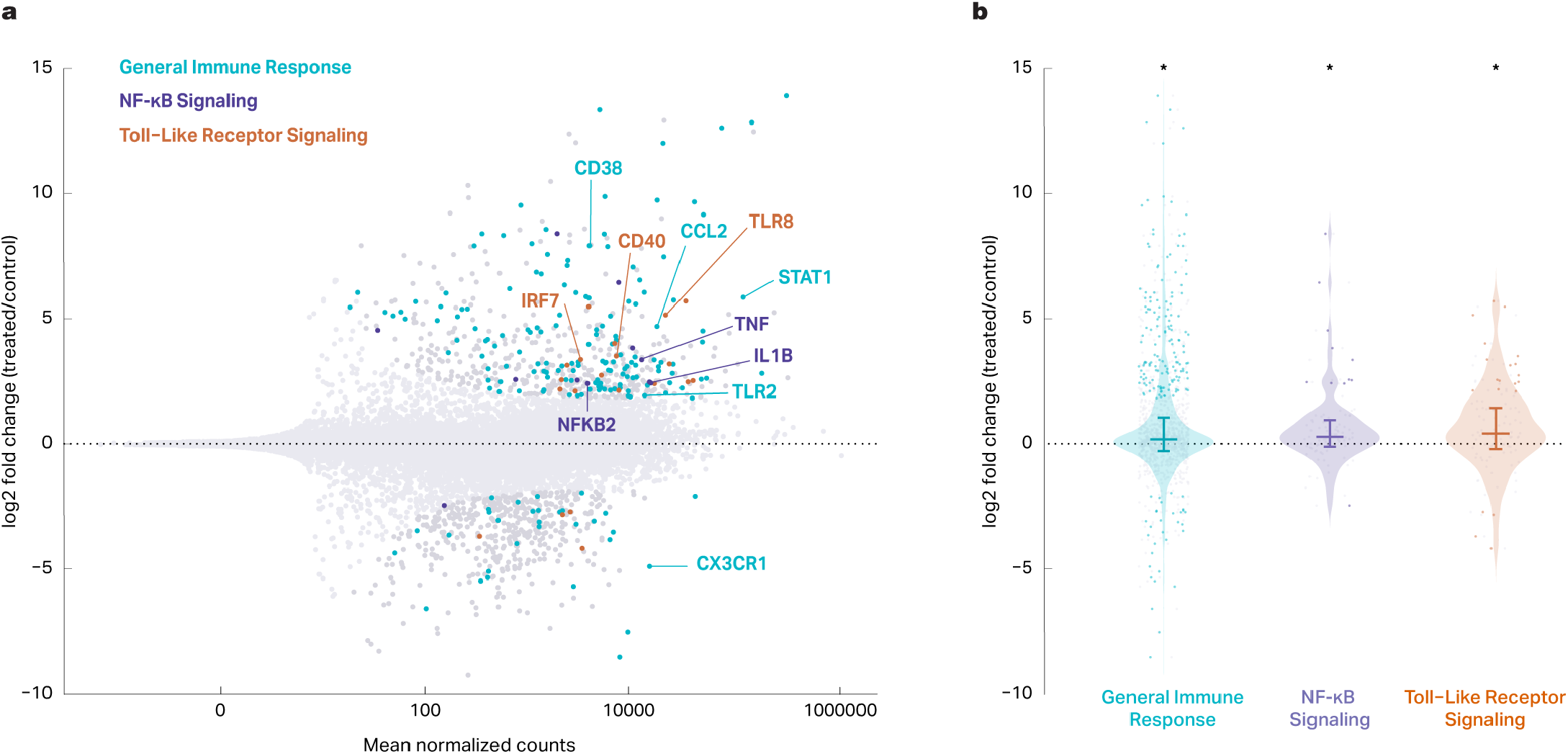
LPS/INFγ treatment induces a robust proinflammatory transcriptional change in IMGLs. **(a)** Bulk RNASeq was performed in IMGLs treated with 1μM Aβ oligomers or 10nM LPS+20nM INFγ for 24h. Differential genes (p< 0.01, absolute fold change > 2) are labeled in dark gray and non-significant genes (p > 0.01, absolute fold change < 2) are labeled in light gray. Blue, purple, and orange represent differential genes associated with general immune response, NF-κB signaling, and toll-like receptor signaling respectively. **(b)** Log_2_ fold-changes for the genes in specific pathways shown in 4A. Non-significant genes are shown in light gray and differential genes are shown in blue, purple, and orange respectively for general immune response, NF-κB signaling, and toll-like receptor signaling. 1 Sample Wilcoxon test showed significant differences for the genes in each pathway vs 0 (p=6.38 × 10^−18^, p=1.28 × 10^−4^, p=1.88 × 10^−4^ respectively).

To assess the validity of LPS/INFγ treated IMGLs as a model of AD, we compared the transcriptional profiles of our LPS/INFγ-treated IMGLs to published DAM gene signatures from AD mouse models^38^. DAM genes were significantly enriched among LPS/INFγ differentially expressed genes (Fisher’s Exact Test, p=4.17 × 10^−15^). The directional effects of our LPS/INFγ-treated IMGLs were also concordant with the directional effects of the DAM signature (**Figure 5**). Genes upregulated in DAM tended to be upregulated in IMGLs upon treatment with LPS/INFγ (median log2 fold-change = 0.26, Wilcoxon test, p=1.46 × 10^−6^), while genes that are downregulated in DAM also tended to be downregulated in IMGLs upon treatment with LPS/ INFγ albeit to a lesser degree (median log2 fold-change = -0.09, Wilcoxon test, p=0.02) (**Figure 5**). Among the shared upregulated genes are MS4A7, which has been shown to regulate TREM2 protein and to be upregulated in late-onset AD brains^39,40^, and ITGAX which has been shown to be upregulated in aged microglia^41,42^. We note that the effect sizes of LPS/INFγ on DAM genes are small; however, these effect sizes are comparable to those observed in the mouse models used to identify DAM genes (median log2 fold-change of 0.17 and -0.23 for upregulated and down-regulated DAM genes respectively). Taken together, these results suggest that LPS/INFγ-treated IMGLs recapitulate many features of the DAM transcriptome.

**Fig. 5.**
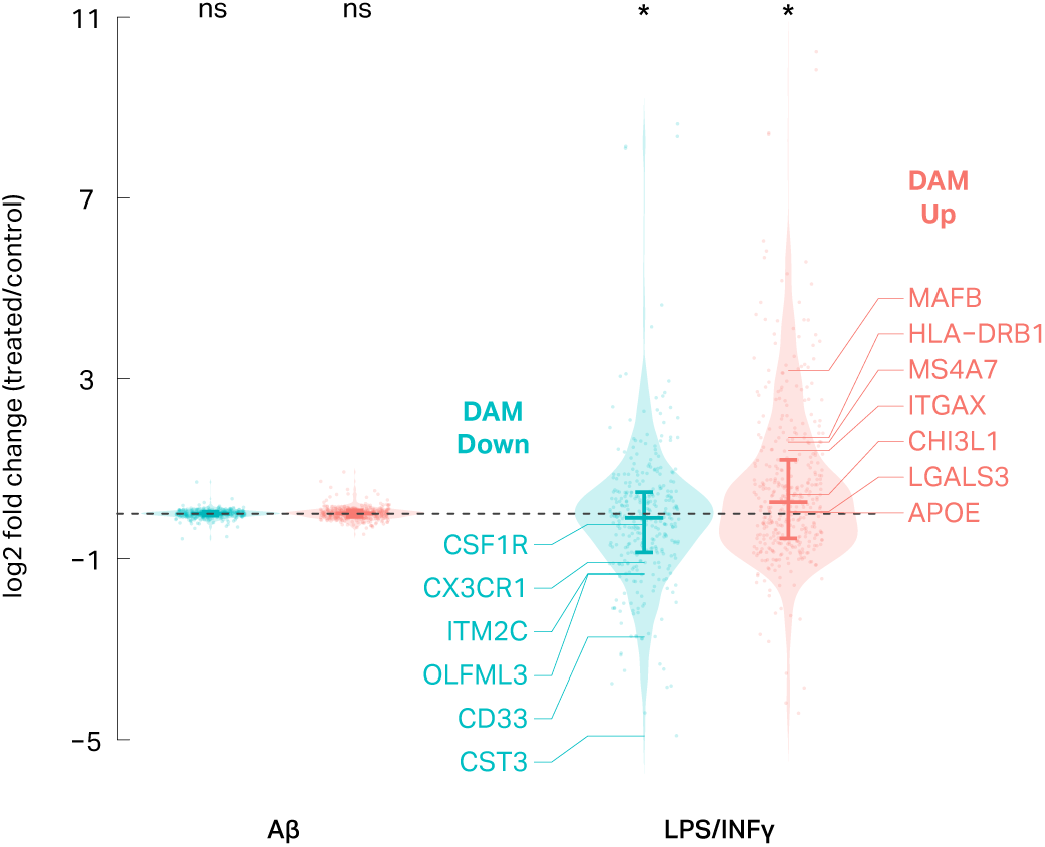
Treatment of IMGLs with LPS/INFγ induces transcriptional changes similar to those observed in disease associated microglia. Violin plots depict how DAM genes change in IMGLs treated with Aβ (left) or LPS + INFγ (right). DAM down and DAM up genes are significantly down and upregulated, respectively, in IMGLs treated with LPS + INFγ (Wilcoxon test p=0.02 and p=1.46 × 10^−6^ respectively).

## Discussion

Progress to unveil the mechanisms behind the origin and progression of AD has been slow in part due to the difficulty in finding adequate models to mimic this disease in the laboratory. Although mouse models have shown to be useful in many aspects, the genetic and physiological differences between species have limited certain areas of research^43^. Conversely, access to human brain samples is limited and the molecular identity and stability of some cell types, mostly brain immune cells, once extracted from the brain have been controversial^44,45^. As a result, the development of human in vitro cell culture models, particularly IPSC-derived celular models, have been broadly used to mimic AD in a dish^46–48^. This alternative has been particularly valuable since it is amenable to genome engineering, disease modeling, and high-throughput screening techniques.

Here, we analyzed the feasibility of using these in vitro cell models, including several established primary and iPSC-derived immune cell lines, and their similarity to human microglia using a genome-wide transcriptomic approach. Not surprisingly, we identified IMGLs as the most similar in vitro culture to mimic human primary microglia (**Figure 1**). This agrees with previous studies done both using RNA-seq^20,24^ or microarray and qPCR approaches with fewer cell types^49,50^.

When the use of IMGLs is not possible due the complexity and high costs of their obtention, our results also suggested that the use of PBMC Mϕ or even macrophages derived from the THP-1 cells are preferable before the use of the HMC3 microglia established cell line, which has fewer similarities with primary microglia. Interestingly, a poor transcriptional similarity between a murine established microglia cell line (BV2) and primary microglia has also been previously reported^51^, raising concerns about the established microglia cell lines that are most often selected to study neuroinflammation in neurodegenerative diseases.

The stimulation of different innate immune cells in culture with synthetic Aβ has become a common approach to model AD in vitro, as Aβ accumulation in the brain is considered one of the main triggers for the development of AD and immune brain cells have been identified as the main players in AD origin and progression. Although the phagocytosis of this component into in vitro microglia-like cell cultures has been previously shown^24,52^, very few studies have shown any kind of pro-inflammatory response after this treatment. And those studies that did investigate inflammation focused on only a handful of proinflammatory genes and proteins quantified via qPCR, micorarrays, or western blots^24,53–55^.

Here we analyzed the effect of synthetic Aβ treatment in innate immune in vitro cell cultures using the same genome-wide transcriptomic approach with a variety of experimental conditions. To our surprise, even though Aβ is phagocytosed inside the cells (**Figure 3B**), we found that it does not trigger a robust transcriptional response in any of the tested cell types or conditions (**Figure 2**).

Several factors could explain why we did not observe robust transcriptional activation in these experiments. First, the synthetic Aβ used here has known differences in structure, heterogeneity and toxicity compared with the Aβ found in human AD brains. Aβ isolated from AD patients comprises a complex mix of Aβ peptides with heterogeneous lengths and posttranslational modifications,^27,56–62^. The use of Aβ preparations extracted from postmortem human brains using previously published extraction protocols^63^ could give more insight into this hypothesis in future studies. Second, the microglia response to Aβ and the consequent neurotoxic effect could be driven uniquely by post transcriptional events, which would agree with previous reports describing the relevance of this layer of regulation in the control of the innate immune response^64–66^ and previous reports of increased proinflammatory cytokine release after Aβ treatment^53,67^. Third, these mono cultured cell lines may not accurately reflect the biology of a human brain which involves a vast array of intertwined cell types. Indeed, studies in animal models have revealed inflammation and microglial activation following injection of synthetic Aβ^68–70^. More work needs to be done to determine if a similar response is observed in brain organoids.

Regardless of these possible explanations, our results suggest that using in vitro immune cell cultures in the presence of synthetic Aβ might not be an adequate and/ or sufficient model to mimic the microglia response in the context of the AD brain. Moreover, this suggests that Aβ phagocytosis alone is not sufficient evidence of microglia activation, at least at the transcriptional level.

Even though these results do not disprove the hypothesis that in the human brain, Aβ induces an exacerbated inflammatory response driven by brain immune cells, and that this plays a major role in the origin and progression of AD, it adds evidence that supports the current concern about placing the Aβ cascade hypothesis as the lone explanation for the AD etiology^71^. More factors, such as the role of tau, microbial infections, and more complex interactions between different brain cell types should be considered and further investigated to understand the origins of the neuroinflammation state observed in the AD brain.

Finally, in agreement with recent studies^33^, we showed that IMGLs exhibit a significant transcriptional response to LPS/INFγ. While this response is smaller than the one observed in THP-1 cells, as was also previously reported^50^, it resembles the transcriptional profile of DAMs in the AD brain. These results suggest that treating IMGLs with LPS/ INFγ may be a suitable platform to study AD-related microglia inflammation.

## Methods

### Amyloid beta formation

Amyloid-beta (Aβ_1-42_) (AnaSpec; Fremont, CA) was prepared using two different methods as indicated in the figure legends (Methods A or B). When not explicitly indicated, Aβ generated following Method A was used. In the Method A Aβ was generated as previously described^23^. Amyloid-beta (Aβ_1-42_) peptide (AnaSpec; Fremont, CA) was dissolved in Hexafluoro-2-propanol (HFIP, Sigma) to 1mM, incubated for 1 hour, then left to evaporate in a vacuum concentrator overnight allowing monomerization. To form oligomerized Aβ (oAβ), the dried film was reconstituted in DMSO (Sigma) to a concentration of 5 mM, and then diluted to 15 μM in PBS (pH 7.4) and stored at room temperature for 30 minutes. To form fibrillarized Aβ (fAβ), the dried film was reconstituted in a 40 mM Tris-HCl (pH 8.0) and 250 mM NaCl solution, incubated at room temperature with agitation for 6 days, then centrifuged for 10 mins at 15000g. The resulting pellet was resuspended in PBS to a concentration of 15 μM. OAβ and fAβ formation were confirmed by western blot as it is described below. Aβ was thoroughly mixed and diluted in media to the desired concentration prior to cell exposure. FAβ contained some oligomers as indicated by western blot analysis.

In method B, Aβ was reconstituted as previously described in Abud, 2017. Briefly, Aβ was dissolved in 75ul of NH4OH (0.1%) then filled to 1 ml with sterile PBS to achieve a 1 mg/ml concentration. Aβ was then diluted to 100 μg/ml with sterile water, vortexed, and incubated at 37°C for 30 mins to form oligomerized Aβ (oAβ), and 7 days or 5 days to form fibrillarized Aβ (fAβ). Aβ was thoroughly mixed and diluted in cell culture media to the desired concentration prior to cell exposure. FAβ contained some oligomers as indicated by western blot analysis (**Figure 2A**).

### Characterization of Aβ using Western blot

The size of the resulting Aβ preparation after performing each protocol was characterized using SDS-PAGE and western blotting. 120 ng of proteins were separated by electrophoresis in a 12% gel and transferred to a nitrocellulose blotting membrane. After blocking, the membranes were incubated overnight at 4°C with anti-Aβ primary antibody (6E10 Biolegend, SIG-39320, 1:1000) following incubation with goat anti-mouse secondary antibody (Thermo Scientific, 1:1000) during 1h. The membranes were subsequently washed and placed in the Radiance Plus Chemiluminescent Substrate (Azure Biosystems, Dublin, CA, USA) and visualized using the Azure c600 gel imaging system (Azure Biosystems).

### Immortalized human cell lines

The immortalized human monocyte cell line THP-1 (ATCC, #:ATCC ® TIB-202TM, obtained from the Tissue culture facility at UNC Chapel Hill), was cultured in RPMI 1640 with L-Glutamine-Bicarbonate (Sigma-Aldrich # R8758) containing 10% FBS (Thermo Fisher Scientific, GibcoTM # 10500064), and 1% penicillin/streptomycin (Sigma-Aldrich #P4333) in an atmosphere of 5% CO_2_ at 37ºC. To differentiate THP-1 monocytes into macrophages, cells were plated at a density of 1.0 × 10^6^ cells/well in 6-well plates and incubated with 25 μM PMA (12-O-tetradecanoylphorbol) during 24 h. Macrophages were then allowed to rest for 72h before Aβ or LPS/INFγ treatment as described below. The HMC3 immortalized microglia cell line (obtained from ATCC # CRL-3304) and the astrocyte-like U-87 cell line (ATCC # 30-2003 obtained from the Tissue culture facility at UNC-Chapel Hill) were both cultured in EMEM (ATCC # 30-2003) containing 10% FBS and 1% penicillin/ streptomycin in an atmosphere of 5% CO_2_ at 37ºC.

Cells were treated with 1μM Aβ or 10nM LPS/20nM INFγ (Sigma-Aldrich # L2630 /Peprotech # 300-02) during 24h. Treatments were carried on in serum-free media with 1% penicillin/streptomycin (100 U/ml penicillin/100 μg/ml streptomycin) unless otherwise stated.

### Generation of peripheral blood mononuclear cell-derived macrophages

Human peripheral blood mononuclear cells (PBMCs) were isolated from deidentified healthy donor blood obtained from the New York Blood Center (Long Island City, NY). Research was approved by the center prior to acquisition. Approximately 10^8^ PBMCs were isolated using Ficoll-Paque density gradient centrifugation (MilliPore Sigma # GE17-5442-02). Cells were cultured in DMEM 10% FBS and seeded in 6-well ultra-low adhesion plastic culture dishes. Monocytes were allowed to adhere for a period of 5-7 days in vitro. Media was then exchanged to DMEM containing 15 ng/mL recombinant human granulocyte-macrophage colony stimulating factor (rhGM-CSF; R&D Systems # 215-GM-010/CF). PBMC Mϕ were allowed to differentiate for an additional 5 days and were returned to DMEM 10% FBS until stimulation.

### Generation of IPSCs-derived microglia

IPSCs-derived microglia cells were generated as previously described (McQuade et al. 2018). Briefly, IPSCs (Wicell UCSD021i-3-9) were differentiated into Hematopoietic progenitor cells (HPCs) using the STEMdiff™ Hematopoietic Kit (Stem Cell Technologies # 05310) according to manufacturer instructions. IPSCs cells were grown in mTeSR1 (STEMCELL technologies Catalog # 85850) and passaged with 0.5 mM EDTA in PBS (-Ca^+2^/Mg^+2^). On day -1 cells were plated into mTeSR1 medium with 0.5 μM Thiazovivin (STEMCELL technologies # 72252) onto hESC-matrigel coated (Corning # 354277) 6-well plates at different densities. On day 0 plates with a density of 10 to 20 aggregates per cm^2^ with a size ranging from 0.1-0.2μm were selected. A small aggregate density and size was found to be critical at this stage. mTeSR1 medium was replaced with medium A (2 mL per well of a 6-well plate). On day 2 50% of the medium A (1mL/well) was removed and replaced with 1 mL of fresh Medium A per well. On day 3, all medium was carefully removed and 2 mL/well medium B was added. Without removing media, cells were supplemented with 1 mL/well of medium B on days 5, 7, 9 adding the media carefully to not disturb the cells. On day 12 non-adherent cells (HPCs) were collected carefully to not disturb adherent cells and centrifuged 300 x G 5 min. The differentiation of HPCs into IMGLs was performed using the STEMdiff™ Microglia Differentiation Kit and the STEMdiff™ Microglia Maturation Kit (STEMCELL technologies # 100-0019 and #100-0020) according to manufacturer instructions. Briefly, HPCs were plated at a density of 1-2 x10^5^ cells per well in hESC-matrigel coated 6-well plates using 2ml of Microglia differentiation media. On day 0 until day 24, 1ml of fresh media was added to each well, except for day 12, in which all the media except for 1ml was collected and spinned down 300 G x 5min, cells were resuspended in 1ml of fresh media and returned to the well. On day 24 cells were collected, centrifuged 300 G x 5min and counted. 1ml of conditioned media per well was left and fresh Microglia Maturation Media media was added to achieve 1×10^6^ cells in 2ml. Cells were plated (2ml/ well) in fresh hESC-matrigel coated 6-well plates. 1ml of fresh media was added every other day until cells were used on days 28-30. On day 28-30 cells were semi-adherent, all media except for 1ml was collected, centrifuged 300 G x 5min and cells were resuspended in 1ml/well of Microglia Maturation media with or without 2x concentration of Aβ or LPS/INFγ and returned to the well. After 24h treatment, cells were collected for RNA extraction.

### Cell labeling and Aβ treatment

For live-cell imaging, 50,000 iPSC-microglia or THP-1 macrophages were cultured in a vitronectin-coated eight-chambered cover glass dish (Cellvis, C8-1.5H-N). THP-1 macrophages were labeled with 2 μM CellTracker Green CMFDA dye (Invitrogen, C2925) in HBSS containing Ca^++^ and Mg^++^ (Gibco, 14025-092) for 30 minutes at 37°C. Medium was replaced with cell medium containing 2 μg/ mL β-Amyloid_1-42_-HiLyte Fluor 555 and cells were imaged 1 hour following treatment.

### Live-cell imaging

Confocal images were acquired on an inverted Zeiss 800 laser scanning confocal microscope equipped with 405, 488, 561, and 647 nm diode lasers, gallium arsenide phosphide (GaAsP) detectors, and a transmitted light photomultiplier tube (PMT). Images were acquired using a 63X/1.4 NA objective lens at 37°C and 5% CO_2_ (Carl Zeiss, Oberkochen, Germany).

### RNA-seq

Immediately after cells were harvested, RNA was extracted using RNeasy Mini Kit (QIAGEN) following manufacturer’s guidelines. RNA integrity number (RIN) was measured for all samples using Agilent tapestation 4150 system. All sequencing libraries analyzed were generated from RNA samples measuring a RIN score ≥ 8.5. RNA-seq libraries were generated using the KAPA RNA HyperPrep with RiboErase kit using 500 ng of isolated mRNA as input and following manufacturer’s instructions. Libraries were quantified and normalized using DNA tapestation and sequenced as paired-end 75-base-pair reads in the Illumina Nextseq 500 platform.

### RNA-seq quantification

RNA-seq libraries were sequenced to an average depth of approximately 55-70 million reads per sample. Low-quality reads and adapters were trimmed using Trim Galore! (v. 0.4.3), and trimmed reads were then mapped to the hg19 transcriptome (GENCODE, release 19) and quantified using Salmon’s mapping-based mode (v. 0.8.2)^72,73^. Both programs were run with default settings, using paired-end inputs. Gene-level quantifications were summarized from each sample using the R package tximport (v. 1.2.0)^74^.

### Principal component analysis and variance partitioning

The untreated controls from each experiment were analyzed separately for the purpose of principal component analysis (PCA), seen in Figure 1, and characterizing drivers of variance. The read counts summarized by tximport were variance-stabilizing transformed (VST) in R using DESeq2 (v. 1.22.2)^75^. Genes were subset for only those with 100 or more transcripts per kilobase million (TPM) in at least one sample (n = 5,192). The VST counts for these genes were used to calculate variance for PC1 and PC2. Variance was characterized using the R package variancePartition (v. 1.22.0) using the same filtered VST counts, and the following formula: ∼ (1|Tech_Rep) + (1|Bio_Rep) + (1|Cell_Type) + (1|Source)^21^. The fitExtractVarPartModel function then fits the VST counts to a linear mixed model and determines the fraction of variance attributable to each covariate in the design formula, with any unexplained variance attributed to residuals. The mean percentage of variation across all genes was reported.

### Differential RNA-seq analysis

Differential analysis was conducted in R with DESeq2 (v. 1.22.2), using a design adjusting for replicate variability when calculating differences between treatment groups (∼ replicate + condition). Each experiment was analyzed separately to identify the number of differential genes caused by each treatment relative to untreated controls. Differential genes were defined as genes with an FDR-adjusted p-value below 0.01 (Wald test, log2 fold-change threshold of 1) and an absolute fold change greater than 2 when comparing treated samples to their respective controls. Fold changes were shrunken using the “apeglm” method within DESeq2 (v. 1.14.0)^76^.

### GO and KEGG Enrichment Analysis

The “findMotifs.pl” tool in the HOMER software suite (v. 4.10.4) was used on genes that were upregulated in IMGLs upon LPS/INFγ-treatment to identify significantly enriched GO terms (p < 0.05) and Kyoto Encyclopedia of Genes and Genomes (KEGG) pathways (p < 0.05)

### DAM Enrichment Analysis

IMGL RNASeq data was subset for genes present in DAM gene signatures (DESeq2; > 100 counts per million). Fisher’s exact test was used to determine the enrichment of DAM genes, using all genes expressed in IMGLs as background (DESeq2; > 10 counts per million). Wilcoxon rank sum test was used to determine whether each group was significantly different from zero.

## Data availability

Raw FASTQ files are available on SRA under accession number PRJNA750801. Transcript-level quantifications from Salmon are available on GEO under series GSE181153, in addition to a single gene-level counts table with each sample represented as a column, as generated by tximport. The GEO series page also includes a table describing the samples used in DESeq2 for each column of Figure 3.

## Acknowledgments

We would like to acknowledge Erika Deoudes for illustrations and graphic design. IYQ is supported by a Bright-Focus Foundation postdoctoral fellowship. Acknowledgement is made to the donors of the Alzheimer’s Disease Research program that supports this fellowship. AEC was supported by the UNC Postbaccalaureate Research Education Program R25GM089569. MLB is supported by the NIH/NIGMS training grant T32GM135128. BAE is supported by NSF-GRFP DGE-1650116. JVR and SC are supported by NIA, R00AG052570. RBM is supported by R01 NS108808. HW, TJC and DHP are supported by NIH grant R01AG066871.

## Supplemental Information

**Supplementary Fig. 1.**
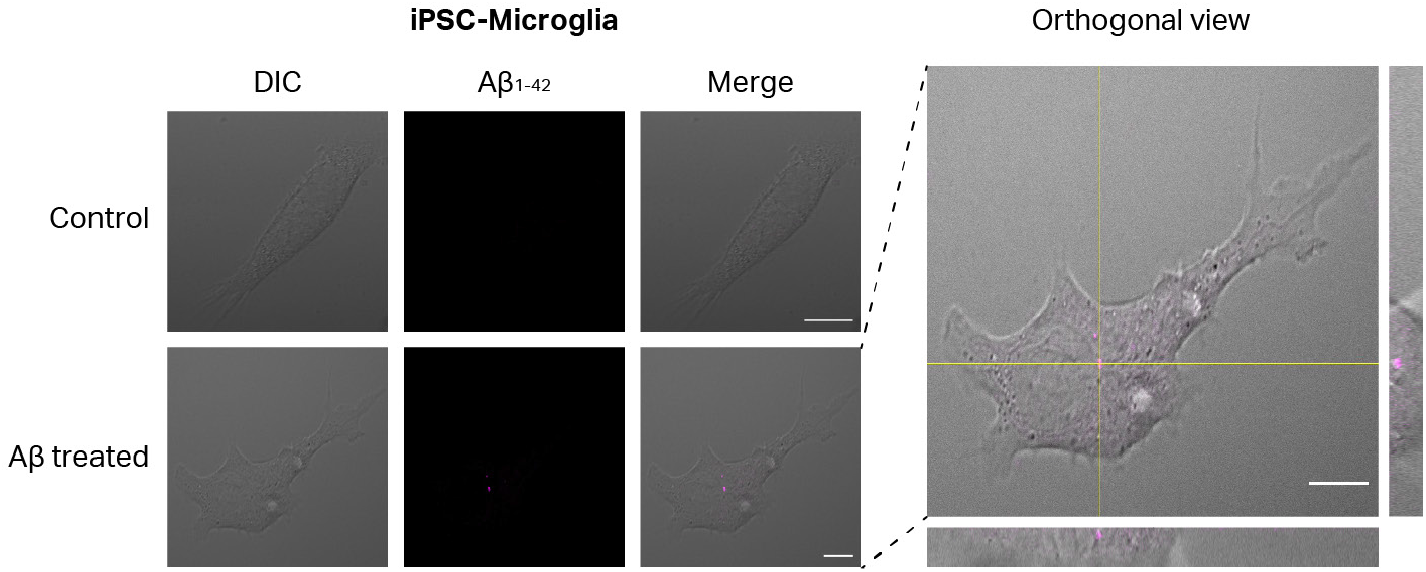
Aβ is incorporated into IMGLs. IMGLs incubated with 2μg/mL of oligomeric β-Amyloid_1-42_-HiLyte Fluor 555 for 1 hr. Confocal images were acquired on an inverted Zeiss 800 laser scanning confocal microscope. Scale bar, 10 μm.

